# Instance-based Transfer Learning Enables Cross-Cohort Early Detection of Colorectal Cancer

**DOI:** 10.1101/2025.02.22.639690

**Authors:** Yangyang Sun, Shunyao Wu, Zhenming Wu, Wenjie Zhu, Hao Gao, Jieqi Xing, Jin Zhao, Xiaoqian Fan, Xiaoquan Su

## Abstract

Colorectal cancer (CRC) continues to be a major global public health challenge. Extensive research has underscored the critical role of the gut microbiome for diagnostics of CRC. However, early-stage prediction of CRC, particularly at the precancerous adenomas (ADA) stage, remains challenging due to the instability of microbial features across cohorts. In this study, we conducted a systematic analysis of 2,053 gut metagenomes from 14 globally-sampled public cohorts and a newly recruited cohort. Despite substantial regional and cohort-level heterogeneity in microbiome composition, we elucidated that the consistent dynamic patterns of microbial signatures are the fundamental for CRC detection. These patterns enabled robust performance in both inter-cohort and independent validations using an optimized bioinformatics framework. In contrast, such basis was lacking in ADA-associated microbial markers, limiting the generalizability of early detection models. To address this, we developed an instance-based transfer learning approach, Meta-iTL, which effectively leveraged knowledge from existing datasets to detect CRC risk at the ADA stage in the newly recruited cohort. Thus, Meta-iTL overcomes challenges posed by cohort-specific variability and limited data availability, advances the application of non-invasive approaches for the early screening and prevention of CRC.

## Introduction

Colorectal cancer (CRC) is one of the most prevalent cancers globally and remains a major cause of cancer-related mortality. According to the American Cancer Society, approximately 106,590 new CRC cases and 53,010 deaths are expected in 2024 alone ^1^. A large number of CRC cases are diagnosed at advanced stages, resulting in poor prognoses and contributing significantly to high mortality rates. Early screening is therefore critical for improving patient outcomes ^2,3^. Traditional screening methods, such as colonoscopy ^4^, are effective for early detection and can reduce CRC incidence and mortality. However, challenges associated with convenience, risks, and the high costs of invasive procedures remain pressing concerns ^4-8^. Consequently, researchers have been actively exploring alternative, non-invasive diagnostic approaches.

As a non-invasive stool-based approach, gut microbiome holds significant potential and has become widely utilized in the detection and study of related diseases ^9-11^. Studies have demonstrated that the gut microbiome plays a pivotal role in the occurrence and progression of CRC ^12^, with microbial markers ^13-15^ identified for predicting disease susceptibility ^16^, progression ^17^, prognosis ^18,19^, and treatment response ^20,21^. However, the dynamic patterns of microbial signatures often differ across cohorts, feature types, marker selection methods, and model training approaches, which hinder the reproducibility and practical application of microbiome-based screening ^22-24^. These limitations underscore the urgent need for systematic, large-scale analyses of gut microbiomes across multiple cohorts and the development of optimized bioinformatics protocols for CRC detection.

On the other hand, it is estimated that 80% of CRC cases arise following the adenoma-carcinoma sequence, which involves the progressive accumulation of mutations in a period of 10 -15 years on average ^25,26^. The early screening of CRC at precancerous-stage adenoma (ADA) has brought the 5-year relative survival rate to around 90% ^27^, significantly facilitating early clinical decision making, alleviating the incidence of CRC, and reducing economic burden ^2,3^. However, research on the association between the microbiome and ADA remains limited. Few studies have examined microbial changes in ADA samples ^3,13,28-30^, and most only focused on single cohorts or specific populations with small sample sizes, restricting the generalizability of microbiome-ADA linkage on a broader population ^28,30^. For example, Thomas et al. ^28^ conducted a meta-analysis of metagenomic data for ADA samples but reported poor performance in distinguishing ADA from healthy controls (e.g., area under the receiver operating characteristic curve (AUROC) < 0.6). Thus, accurate detection of ADA across diverse cohorts remains a major challenge.

In this study, we performed a systematic and comprehensive analysis of over 2,000 gut metagenomes from diverse regions worldwide. Despite substantial regional and cohort-level heterogeneity in microbiome composition, we revealed that consistent dynamic patterns of microbial signatures play a fundamental role in cross-cohort CRC detection. Oppositely, such basis was lacking in ADA-associated microbes, challenging the prediction of CRC risk at the early stage. Here we developed Meta-iTL, an instance-based transfer learning (TL) modeling strategy. By leveraging knowledge transfer between samples to be tested (i.e., target domain) and existing data repository (i.e., source domain), Meta-iTL overcomes limitations of the cohort-specific effects and scarcity of ADA cases for training, improving cross-cohort applicability of data models. This approach advances the development and application of non-invasive early screening, offering significant potential for improving early diagnosis and prevention of CRC.

## Results

### Overview of CRC detection and prediction using multi-cohort gut metagenomes

To address the challenges posed by data heterogeneity and bioinformatic variability in linking the gut microbiome to CRC, we conducted a systematic and comprehensive analysis of 2,053 human gut paired-end sequenced metagenomes (**Fig.1a**; **Table 1**). These samples were derived from 14 published cohorts and one in-house recruited cohort (i.e., CHN_WF in **Table 1**; refer to **Materials and methods** for details), spanning three health statuses: 735 samples from CRC patients, 349 from individuals with ADA, and 969 from healthy controls (CTR). Samples exhibited high geographical diversity that surveyed from ten countries/regions and were generated using different sampling procedures, storage conditions, and DNA extraction protocols ^31,32^ (**Table S1**). Importantly, all samples were collected prior to any therapeutic interventions to avoid treatment-related biases ^21,33,34^. Raw metagenomic sequences were filtered by quality control and host removal, and then taxonomical and functional profiles were parsed by MetaPhlan4 ^35^ and HUMAnN3 ^36^, respectively, with consistent parameters setting across all datasets (refer to **Materials and methods** for details).

**Table 1.** Metagenome cohorts and samples.

(a) CRC discovery dataset.
| Cohort | # of CTR | # of CRC | # of ADA | Country/Region | Source |
| --- | --- | --- | --- | --- | --- |
| AT<sup>13</sup> | 58 | 46 | 46 | Austria | PRJEB7774 |
| IND<sup>38</sup> | 30 | 30 | 0 | India | PRJNA531273<br>PRJNA397112 |
| ITA<sup>28</sup> | 51 | 59 | 27 | Italy | SRP136711 |
| JPN<sup>15</sup> | 245 | 111 | 67 | Japan | DRA008156 |
| GE<sup>31</sup> | 57 | 20 | 0 | Germany | PRJEB27928 |
| CHN_HK<sup>39</sup> | 53 | 75 | 0 | HK, China | PRJEB10878 |
| CHN_SH<sup>40</sup> | 85 | 80 | 0 | China | PRJNA731589 |
| FR<sup>41</sup> | 66 | 88 | 40 | France & Germany | ERP005534 |
| US<sup>42</sup> | 48 | 46 | 0 | USA | PRJEB12449 |
| CHN_WF | 48 | 48 | 54 | China | PRJNA1198725<br>(in- house<br>produced) |

(b) CRC validation dataset
| Cohort | # of CTR | # of CRC | # of ADA | Country/Region | Source |
| --- | --- | --- | --- | --- | --- |
| CHN_SH-2<sup>43</sup> | 44 | 32 | 23 | China | PRJNA514108 |
| CHN_SH-3<sup>44</sup> | 100 | 100 | 0 | China | PRJNA763023 |

(c) Additional ADA dataset
| Cohort | # of CTR | # of CRC | # of ADA | Country/Region | Source |
| --- | --- | --- | --- | --- | --- |
| US-2<sup>45</sup> | 28 | 0 | 26 | USA & Canada | PRJNA389927 |
| US-3<sup>46</sup> | 9 | 0 | 26 | USA | PRJNA745329 |
| CHN_SH-4<sup>47</sup> | 47 | 0 | 40 | China | PRJNA706060 |

For our exploratory analysis of CRC, we designated 10 cohorts as the ‘discovery’ dataset (**Table 1a**), focusing on distinguishing samples with CRC from healthy controls. To comprehensively study on the feasibility of cross-cohort CRC detection, we systematically assessed the effects of confounding factors, annotation types, feature selection methods and models, and established an optimal workflow for CRC detection (**Fig. 1b** and **Fig. 1c**; refer to **Materials and methods** for details). The robustness and applicability of the proposed workflow were confirmed by both cross-validation within the discovery dataset and an independent ‘validation’ dataset (**Table 1b**) comprising two external cohorts. However, when applied to ADA detection, the model exhibited markedly limited cross-cohort generalizability (**Fig. 1c**). Here, we developed an instance-level transfer learning framework, Meta-iTL, enabling the pre-diagnosis of ADA in a newly recruited cohort (**Fig. 1c**).

**Fig. 1.**
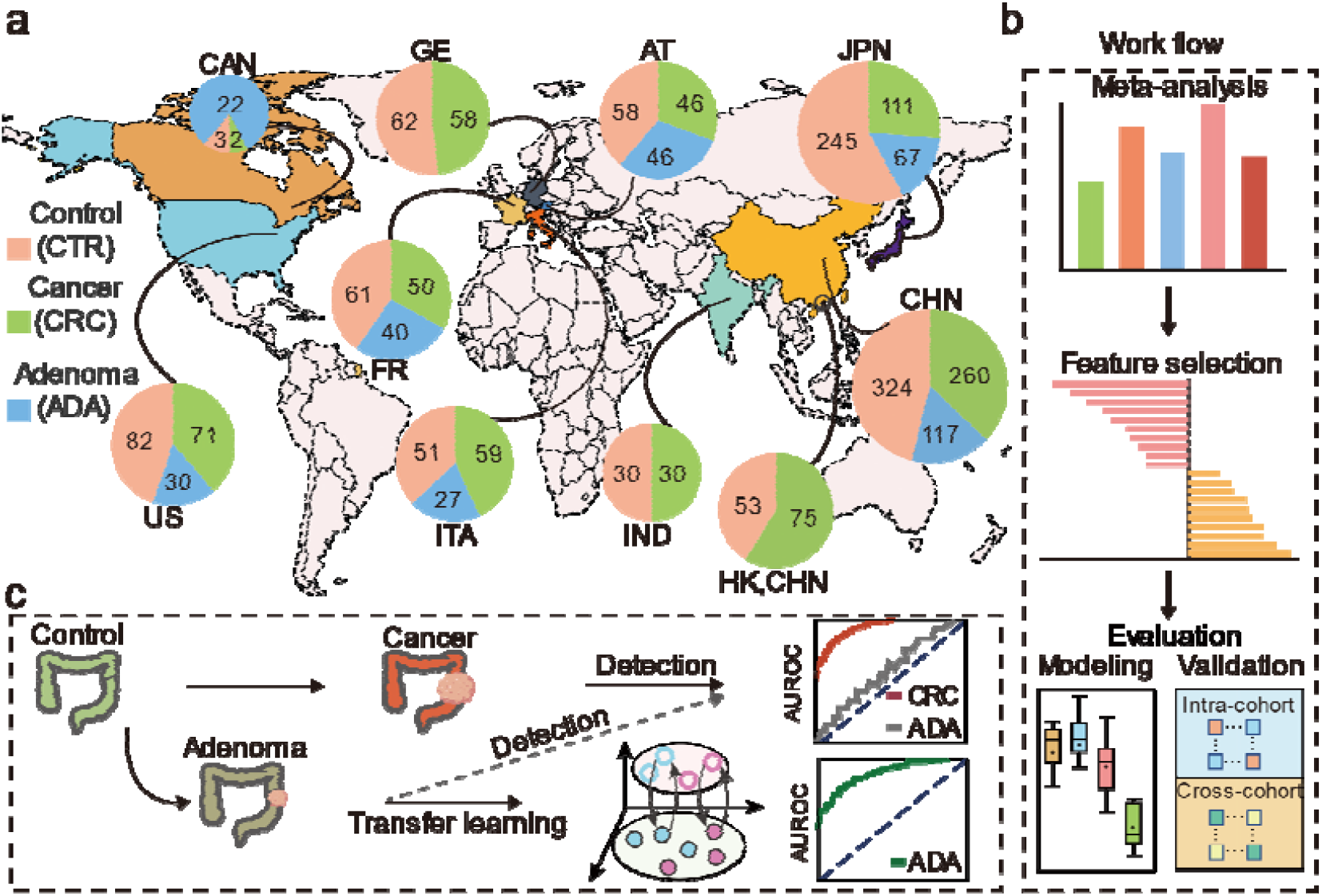
Scheme of cross-cohort integrative analysis of gut metagenomes for CRC early detection. (a) Overview of sample collection and distribution across cohorts. The world map was generated by the Python GeoPandas package (version 0.14.0) ^37^. (b) Analytical framework encompassing effect size estimation of contributing factors, feature selection, data modeling and performance evaluation. (c) Transfer learning enables the cross-cohort detection of CRC at the early stage.

### Impact of confounding factors on microbial composition and mitigation of cohort-related variability

Given the substantial biological and technical differences ^31,48^ among 10 cohorts of the discovery dataset, we investigated the effects of confounding factors on microbial composition and distribution, including cohort-specific variables, BMI (body mass index), age, gender, and health status. The overall beta diversity was measured across eight feature types (taxonomic profiles on class, order, family, genus, species, and SGB (species-level genome bins) ^35^ levels, as well as functional profiles on KO (KEGG Orthology) ^49^ and uniref family ^50^ levels (**Fig. 2a**). Although variance of each microbial component indicated that both cohort and disease status significantly influenced the microbial communities (**Fig. S1**), we found that the cohort-related factor exhibited the largest effect size (**Fig. 2b, Fig. S2** and **Fig. S3**). Despite a shared core of dominant taxa was consistently present across multiple cohorts such as *Ruminococcus_bromii, Eubacterium_rectale, Bacteroides_uniformis, Prevotella_copri_clade_A, Faecalibacterium_prausnitzii* and *Phocaeicola_vulgatus* exceeded 2% relative abundance (**Fig. 2c)**, unique microbial enrichments were evident within individual cohorts. For instance, the IND cohort was characterized by *GGB1627_SGB2230, Prevotella_SGB1680, Prevotella_copri_clade_B*, and *Prevotella_copri_clade_C*, while the ITA dataset showed a predominance of *Alistipes_onderdonkii* and *Alistipes_putredinis*. The AT cohort was enriched with *Anaerostipes_hadrus, Bifidobacterium_adolescentis, Blautia_wexlerae*, and *Candidatus_Cibionibacter_quicibialis*. Additionally, *Escherichia_coli* displayed notable abundance in the AT, CHN_HK, CHN_SH, and CHN_WF cohorts (**Fig. 2c**).

**Fig. 2.**
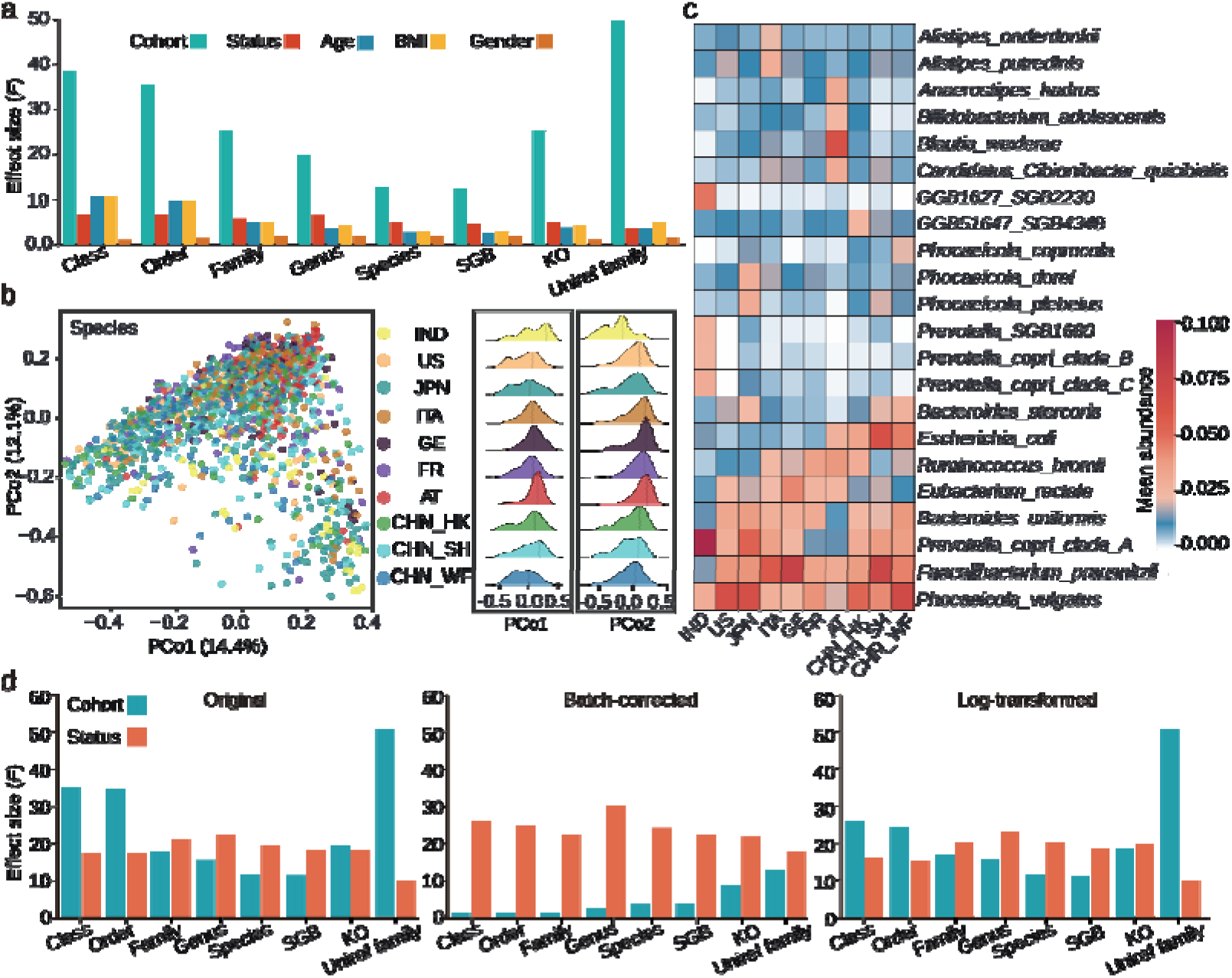
Impacts of different confounding factors on gut microbial composition. (a) The effect sizes of health status and confounding factors on microbial profiles. Each bar represents the PERMANOVA *F-value* derived from microbial features across eight feature types. (b) Beta diversity among discovery dataset cohorts, visualized using Bray-Curtis distances. A series of density plots illustrating the distributions of the first two axes of variation determined by the ordination analysis displayed in principal coordinates analysis (PCoA). (c) Distribution of dominant bacterial communities in each cohort (relative abundance > 2% in at least one cohort). Detailed influences of cohort and health status on eight types of microbial features. Source data is available in Dataset S1.

To explore the influence of technical batch effects, we compared metagenomes sampled from the same city of China conducted by three studies (CHN_SH, CHN_SH-2, CHN_SH-3; **Table 1**). Despite sharing similar geographic origin and health status, significant differences in beta diversity persisted across these cohorts (PERMANOVA test, *p*-value < 0.01; **Fig. S4**). This indicates that technical batch effects ^32,51^, in addition to regional differences ^22^, contributed substantially to microbial variability. Hence, we applied batch correction ^52^ and logarithmic transformation to the relative abundance of profiles (refer to **Materials and methods** for details), respectively. With batch correction, the influence of cohort factors was significantly reduced, which became lower than that of disease (**Fig. S5**). However, such tools should be used with care because they can mistakenly detect and remove actual biological signals from the data ^53^. On the other hand, logarithmic transformation, while less effective at eliminating cohort effects, improved disease status discrimination as annotation resolution was refined (**Fig. S5**). This adjustment balanced the feasibility of detecting CRC across cohorts with the preservation of biological relevance.

Using the three preprocessing strategies (i.e., original, batch-corrected, and log-transformed profiles), we identified microbial features significantly associated with CRC as candidate biomarkers using MaAsLin2 ^54^ (**Table 2**; refer to **Materials and methods** for details). Notably, as the profiling resolution increased (e.g., taxonomy annotation from class level to SGB level), the influence of disease status on biomarkers became more pronounced, surpassing the effect of cohort-related factors. Specifically, at the genus, species, and SGB levels, the PERMANOVA *F*-values for disease status grouping exceeded those of cohort effects (**Fig. 2d**). This finding highlights the importance of selecting high-resolution microbial signatures to minimize cohort and batch variability, thereby focusing on variations directly linked to CRC status across multiple cohorts.

**Table 2.** Feature selection methods.

| Method | Version / Implementation | Distribution | Criteria |
| --- | --- | --- | --- |
| *ANCOM-II*<sup>63</sup> | ANCOM-II function (version 2.1)<br>in R | Non-parametric | W-score>0.8 |
| *MetagenomeSeq*<sup>64</sup> | metagenomeSeq (version 1.36.0)<br>package in R | Zero-inflated (log-)<br>Normal | $p$-value <0.01 |
| T-test | Stats (version 4.1.3) package in R | Normal (default) | $p$-value <0.01 |
| *Wilcoxon test* | Stats (version 4.1.3) package in R | Non-parametric | $p$-value <0.01 |
| *MaAsLin2*<sup>54</sup> | MaAsLin2 (version 1.8.0) package<br>in R | Normal (default) | $p$-value <0.01 |
| *RFECV*<sup>65</sup> | RFECV function in the sklearn<br>(version 1.3.2) library | step=10, cv=5 | Highest accuracy |
| LEfSe<sup>66</sup> | Microeco package (version 1.4.0)<br>in R | Non-parametric | $p$-value <0.05 &<br>LDA>2 |

### An optimal workflow reveals the key microbial patterns for CRC detection

We developed a synergistic feature selection strategy that integrates multiple analytical methods to identify the most informative microbial features for CRC detection (refer to **Materials and methods** for details; **Fig. S6**). To determine the optimal modeling approach (**Fig. 3a**), we further systematically evaluated three preprocessing strategies (raw, log-transformed, and batch-corrected profiles), nine feature selection methods (including 7 existing tools in **Table 2**, our synergistic strategy, and no feature selection), and five widely used machine learning and deep learning algorithms (support vector machine (SVM), multilayer perceptron (MLP), k-nearest neighbors (KNN), Random Forest (RF), and XGBoost). CRC data models were constructed using eight feature types (**Fig. 2a**). This comprehensive benchmarking across the ten discovery cohorts yielded 1,080 analytical combinations (**Fig. S7**). Based on the performance results (**Fig. S7-Fig. S10**; refer to **Materials and methods** for details), we established an optimized bioinformatics workflow for microbiome-based CRC detection (**Fig. 3a**). This workflow demonstrated robust performance, achieving an average AUROC of 0.86 in five-fold cross-validation using RF at the species level, with similarly strong results at the genus and SGB levels (AUROC = 0.85; **Fig. S11**).

**Fig. 3.**
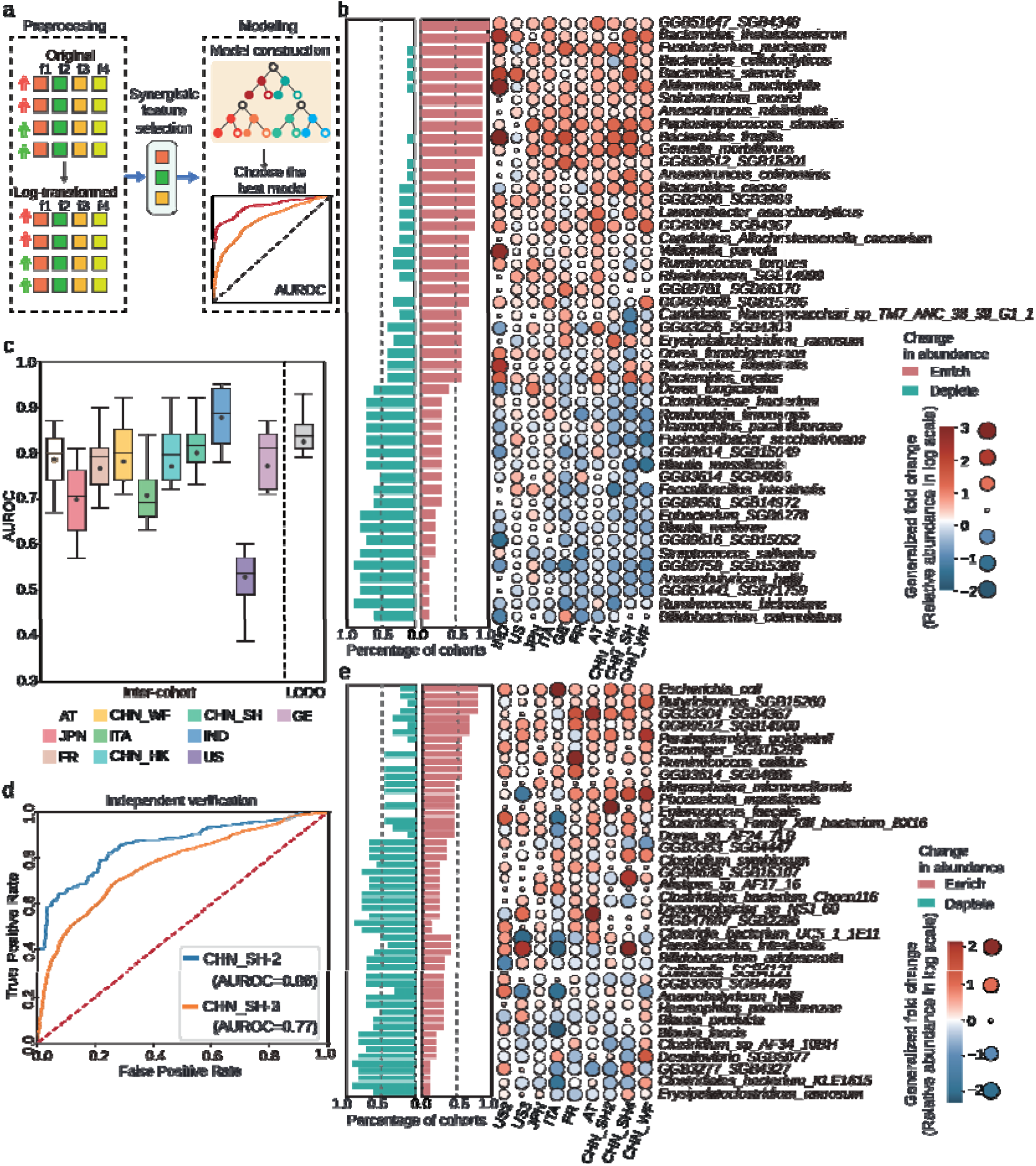
Systematic analysis of cross-cohort CRC detection. (a) The optimal metagenomic bioinformatics workflow for CRC detection. (b) Dynamics of CRC microbial markers across discovery cohorts. Bar heights show the proportion of cohorts with enrichment (red) or depletion (green), and bubbles represent corresponding generalized fold change (gFC) values, with red indicating enrichment and blue indicating depletion. (c) Results of the inter-cohort validation and LODO using profiling on species levels. Each box of the inter-cohort validation summarizes the AUROC values of a single cohort tested by models trained from other individual cohorts separately. The LODO box summarizes the AUROC value of a single cohort tested by models trained from all other cohorts. (d) ROCs of two independent validation cohorts using the model derived from the discovery dataset. (e) Dynamics of ADA microbial markers across multiple cohorts. Source data is available in Dataset S1.

We next performed a detailed examination of the microbial patterns across the ten discovery cohorts at the species level using the above optimal workflow. Results revealed that 82% of the biomarkers identified through synergistic feature selection exhibited consistent CRC-associated dynamics in at least six cohorts (**Fig. 3b**). These species were predominantly affiliated with major phyla, including *Firmicutes, Bacteroidetes, Proteobacteria, Actinobacteria, Fusobacteria, Candidatus_Saccharibacteria*, and *Verrucomicrobia* (detailed listed in **Dataset S1**), mirroring findings reported in previous studies ^41,48^. Specifically, comparative analyses between healthy controls and CRC patients identified 18 species that consistently showed reduced abundance in CRC samples across at least 60% of cohorts. These species were primarily associated with *Firmicutes, Proteobacteria*, and *Actinobacteria*. In contrast, 29 species were consistently enriched in CRC samples, spanning *Firmicutes, Bacteroidetes, Fusobacteria, Verrucomicrobia, Proteobacteria*, and *Candidatus_saccharibacteria*. These findings demonstrated that the microbial signatures identified through the optimal workflow show high consistency in their dynamic changes across health states in multiple heterogeneous cohorts, providing a solid foundation for the development of a reliable, cross-cohort CRC detection model.

Using the optimal workflow and biomarkers screened from all ten discovery cohorts, we further performed inter-cohort validation (training on a single cohort and testing on others) and leave-one-out-of-dataset validation (LODO; make a cohort the testing set and others the training set). Despite the limited sample size in individual cohorts, the average AUROC for inter-cohort validation using the RF model remained at a high level (**Fig. 3c**). We noticed that tests on the US cohort exhibited poor performance, mainly due to the fecal samples in this cohort were freeze-dried prior to DNA extraction and sequencing and stored at -80°C for over 25 years ^42^. After excluding the US cohort as an outlier, the average AUROC of the inter-cohort validation reached 0.79, at the species levels, respectively. LODO validation further improved the AUROC values to 0.85 at the species level (**Fig. 3c**), emphasizing the importance of integrating multi-cohort data for enhanced performance.

The robustness of the workflow across multiple cohorts were further evaluated using two independent validation cohorts from China (CHN_SH-2 and CHN_SH-3; **Table 1b**). The CHN_SH-2 cohort comprised 44 CTR samples and 32 CRC patients, while the CHN-SH-3 cohort included 100 CTR and 100 CRC samples. Using species-level model trained from the ten discovery cohorts, the proposed workflow achieved AUROC values of 0.86 and 0.77 for the CHN_SH-2 and CHN_SH-3 cohorts, respectively (**Fig. 3d**). These results validated the universality of the proposed workflow across multiple cohorts, underscoring the efficacy of using intestinal microbial signatures for accurate and scalable CRC screening.

### Transfer learning enables cross-cohort CRC prediction at precancerous stage

Surprisingly, when applying the optimized bioinformatics workflow, only 50% of ADA-associated asignatures exhibited consistent dynamic patterns across all cohorts (**Fig. 3e**). As a result, a model trained on all other cohorts achieved an AUROC of just 0.60 when tested on the in-house CHN_WF cohort—significantly lower than the 0.86 AUROC observed for CRC detection (**Fig. S12**). This pronounced heterogeneity undermines the effectiveness of cross-cohort models, making within-cohort models the most reliable approach for ADA detection. However, such a strategy can also be constrained by the limited sample sizes available within individual cohorts. In such cases, adopting well-validated models from other locations presents a practical alternative, although careful consideration of model applicability and compatibility is essential for successful implementation.

Here we propose Meta-iTL, an instance-based transfer learning (TL) approach designed to enhance cross-cohort ADA detection (**Fig. 4a**). Basically, Meta-iTL operates by a small set of samples (e.g. *n*=25) from the target domain to capture its specific characteristics, and integrating this information into the source domain of a much larger dataset for model training (**Fig. 4a**, refer to **Materials and methods** for details). Thus, Meta-iTL effectively reduces distributional discrepancies between the source and target domains. This approach addresses two key challenges in ADA detection: the inconsistent behavior of ADA-associated microbial markers across cohorts (**Fig. 4b** and **Fig. 4c**), and the limited sample sizes often found within individual cohorts. Using the CHN_WF cohort as the target domain and all other ADA cohorts as the source domain, we evaluated Meta-iTL for ADA detection. The proportion of CHN_WF samples for TL ranged from 20% to 60%, and 30% samples were randomly selected from the remaining for testing. Each iteration was repeated 50 times to minimize random bias, and both TL-based and non-TL models shared the exact same testing set. Additionally, we performed five-fold cross-validation within the CHN_WF cohort to make a reference AUROC of 0.70 (**Fig. S13a**).

**Fig. 4.**
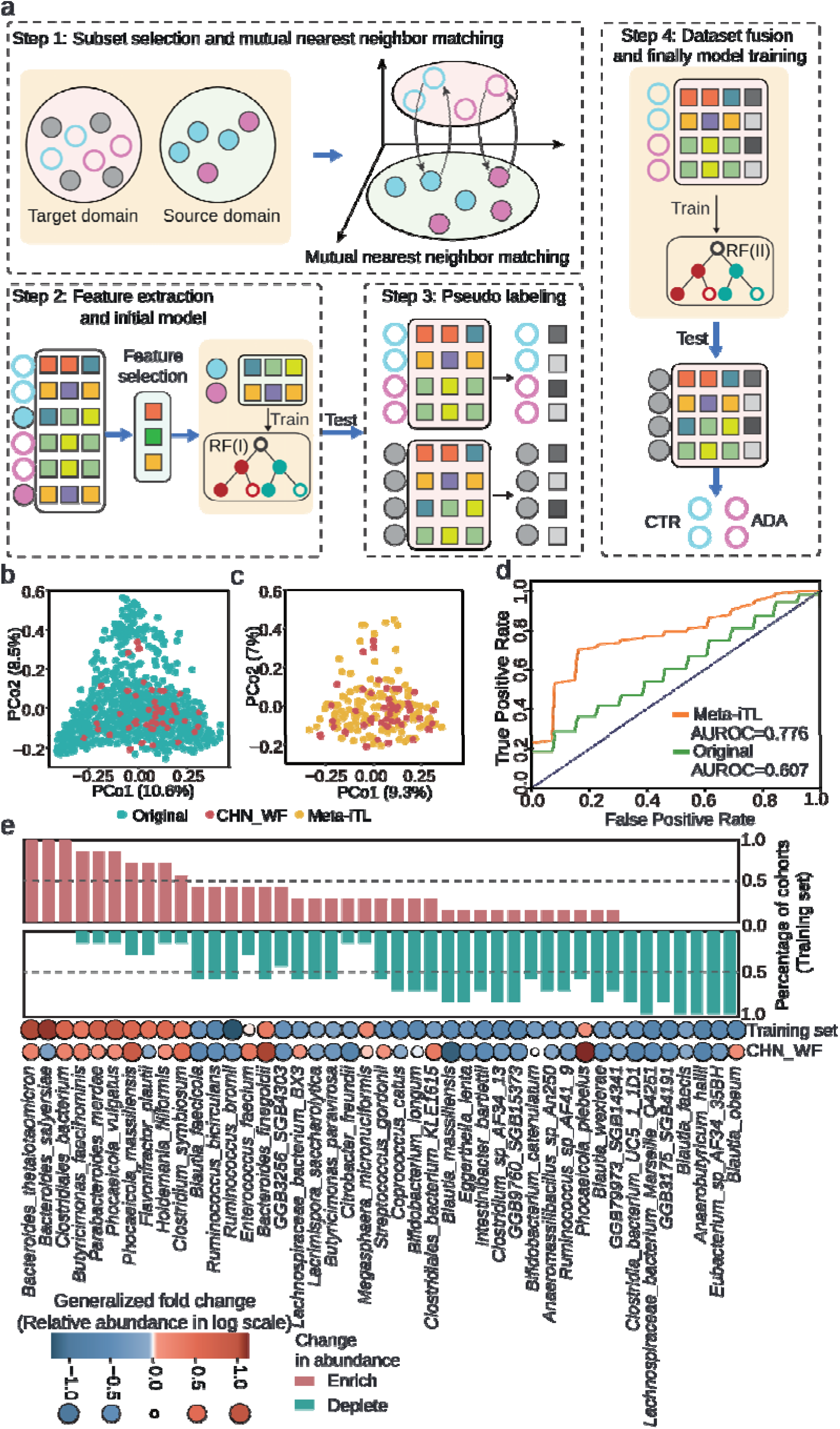
Meta-iTL significantly improves the performance for ADA detection across cohorts. (a) Diagram of the Meta-iTL for instance-based transfer learning. Detailed steps are described in **Materials and methods**. (b) and (c) Principal coordinates analysis (PCoA) based on Bray-Curtis distances between the target domain (CHN_WF cohort) and training sets before and after transfer learning, respectively. (d) ROC curves for ADA detection in the target domain (CHN_WF cohort) before and after transfer learning. (e) Dynamics of ADA microbial markers after transfer learning. Compared to the pattern before transfer learning (**Fig. 3e**), signatures exhibit consistent directional changes within the training sets, while also showed a high concordance rate to the targeted domain (CHN_WF cohort). Source data is available in Dataset S1.

As expected, the source domain model without TL performed sub-optimally in detecting ADA within the CHN_WF cohort that was far below the reference (AUROC=0.61; **Fig. 4d**). When only 20% of the CHN_WF cohort was used for TL, the model’s accuracy did not improve due to insufficient target cohort information. As the proportion of fused CHN_WF samples increased to 30% (*n*=25), the TL model outperformed the non-TL model (**Fig. S13a**). Performance peaked at 40% (*n*=34) target domain integration, achieving an AUROC of 0.78—surpassing the referenced AUROC of the within-cohort model (**Fig. 4d** and **Fig. S13a**). Interestingly, AUROC declined with further increases in target domain sample proportions (e.g., 50% and 60%), likely due to overfitting or reduced influence of source domain data (**Fig. S13a**).

The analysis of microbial composition further elucidated the advantages of the transfer learning strategy. By enhancing the consistency of biomarker dynamics across cohorts, Meta-iTL enabled 66% of candidate markers to exhibit consistent changes in at least five cohorts in the training model. More importantly, this pattern also showed a high concordance rate of 82% with the targeted CHN_WF cohort (**Fig. 4e**). A Jensen-Shannon (JS) divergence analysis also provided insights into Meta-iTL’s efficacy (refer to **Materials and methods** for details). For ADA metagenomes, significant JS divergence was observed between the target and source domains, reflecting substantial differences in their data distributions. After applying Meta-iTL, the JS divergence was notably reduced, indicating that Meta-iTL successfully mitigates gaps caused by cohort-specific factors (**Fig. S13b** and **Fig. S14b)**.

We further validated Meta-iTL by designating the CHN_SH-4 (**Table 1c**) cohort as the target domain. With only *n*=26 samples from the target domain for transfer learning, Meta-iTL achieved an average AUROC of 0.86 (**Fig. S14a**), outperforming the source domain model without TL. Therefore, by integrating a small amount of targeted samples with large-scale data, Meta-iTL holds great promise for the early detection and prevention of CRC by addressing the variability inherent in multi-cohort ADA microbiome datasets.

## Discussions

In this study, we established an optimized bioinformatics framework for cross-cohort CRC detection using one of the largest and most diverse datasets of gut metagenomic samples, encompassing 2,053 samples from 15 cohorts across ten countries and regions. Despite the geographical and technical diversity inherent to metagenomic data across datasets, our synergistic feature selection strategy identified highly consistent dynamic patterns of gut microbial shifts associated with CRC. These findings enabled us to establish a robust workflow capable of achieving high predictive performance across diverse cohorts, highlighting its translational potential for clinical applications. This approach not only enhances generalizability but also ensures that identified biomarkers and models are resilient to data variability. This reproducible framework is readily extendable to other diseases, thereby maximizing the utility of existing large-scale microbiome datasets for diagnostic research and development.

However, substantial heterogeneity in microbial composition across cohorts remains a significant challenge, particularly for early-stage detection. This variability was especially pronounced for adenoma (ADA)-associated biomarkers, which exhibited greater cohort-specific effects compared to CRC biomarkers. Such inconsistencies hindered the accuracy of cross-cohort ADA detection models, posing a critical bottleneck in early CRC screening efforts. To address this, we developed Meta-iTL, an instance-based transfer learning (TL) approach that incorporates a small subset of target domain samples into models trained on large-scale source domain data. By effectively mitigating biases introduced by cohort-specific variability and addressing the scarcity of ADA cases, Meta-iTL significantly improved ADA detection accuracy, offering a practical and scalable solution for enhancing microbiome-based early screening tools in real-world settings.

In conclusion, this study establishes a comprehensive roadmap for metagenome-based CRC early detection and prevention. By providing a standardized bioinformatics workflow and introducing advanced methodologies like Meta-iTL, we aim to bridge the gap between microbiome research and its clinical translation. These findings lay the foundation for developing non-invasive, scalable, and accurate microbiome-based screening tools for CRC and its precancerous stages. As microbiome research continues to advance, our study underscores the critical importance of harmonizing analytical approaches and leveraging innovative machine learning techniques to fully realize the diagnostic potential of the gut microbiome.

## Materials and methods

### Public dataset collection

We used PubMed to search for studies that published fecal shotgun metagenomic data of human colorectal cancer (CRC) patients, adenoma (ADA) patients, and healthy controls (CTR). Raw FASTQ files were downloaded using the SRA toolkit (V.2.9.1) from Sequence Read Archive (SRA) and European Nucleotide Archive (ENA) using identifiers listed in **Table 1**. All metadata for metagenomics were manually curated from the published papers (**Table S1)**.

### In-house fecal sample collection and shotgun metagenomic sequencing

*Subject recruitment and sample collection*. The CHN_WF cohort (**Table 1a**) is an in-house produced dataset. Stool samples were collected from *n*=150 subjects at Shouguang Traditional Chinese Medicine Hospital, Shandong, China. All participants provided written informed consent. Patients with hereditary CRC syndrome or previous history of CRC were excluded from the study. Based on pathology and colonoscopy results, subjects were classified into three groups: *i*. healthy subjects with negative colonoscopy results for tumors, adenomas, or other diseases (CTR; *n*=48); *ii*. subjects with colorectal adenoma (ADA; *n*=54); and *iii*. patients with newly diagnosed colorectal cancer (CRC; *n*=48). The clinical characteristics of the study participants are shown in Supplementary Data1. Fecal samples were collected using sterile specimen collectors and stored at -80°C prior to microbial analysis. This study was approved by the Medical Ethics Committee of Shouguang Traditional Chinese Medicine Hospital (approval no. szy2021-01).

*Metagenomic sequencing*. The DNA from stool samples was extracted using MagPure Stool DNA KF Kit B (MAGEN, Guangzhou, China) according to manual instructions. During the sequencing process, we utilized a short-insert library type and conducted sequencing using the DNBSEQ-T7 platform. The read length was configured to PE150 (paired-end, 150 base pairs each) to ensure comprehensive coverage and high resolution of genomic features. The quality of the raw fastq files was assessed using a Phred+64 encoding system, facilitating accurate measurement of base call quality across the dataset. Raw data were filtered using SOAPnuke ^55^, which involved removing contaminants, adapters, and low-quality sequences. The specific filtering parameters set in SOAPnuke were: “-n 0.01 -l 20 -q 0.4 --adaMis 3 --outQualSys 1 --minReadLen 150”. A total number of approximately 1800 GB of high-quality clean sequence data was obtained.

### Sequence preprocessing and profiling

For all metagenomes, low-quality reads were removed using the Fastp ^56^ tool with parameters “-W 30 -M 20 -y”. The pre-filtered reads were then aligned to the human genome (hg19) using Bowtie2 ^57^ with the parameters “--end-to-end, --no-mixed, --no-discordant, --no-unal, and –sensitive” for host removal. In the profiling step, taxonomic profiles were generated using MetaPhlAn v.4.0.6 (https://github.com/biobakery/MetaPhlAn) with the reference database mpa_vOct22_CHOCOPhlAnSGB_202212 and parameters: “-t rel_ab_w_read_stats and --ignore_archaea”. Functional profiling was conducted using HUMAnN v.3.7 (https://github.com/biobakery/humann) with reference database UniRef90s (version 201901b) and default parameters. The resulting abundance profiles are further summarized into other gene groups, such as KOs (KEGG Orthologs), by using the command “human_regroup_table” with the parameter “--groups uniref90_ko”.

### Preprocessing of microbial features

Microbial taxonomic and functional profiles were first normalized to relative abundances to account for library size. To reduce data noise and enhance data quality, we dropped low-abundance features. At the taxonomic level, features with an average relative abundance of more than 1×10□□ in at least two discovery cohorts were retained, while features with a relative abundance of zero in 90% of the samples were removed, and the standard deviation of the selected features was ensured to be greater than zero. At the KEGG Orthology (KO) level, features with a mean relative abundance greater than 1 × 10□□ across at least eight discovery cohorts were retained, following the same criteria for prevalence and variability. Similarly, at the uniref family level, features with a mean relative abundance exceeding 1 × 10□□ in at least five discovery cohorts were selected, adhering to the same filtering thresholds. The number of features on each type for analysis was summarized in **Table S2**. We then further filtered the data and only retained samples containing at least 50 species for subsequent analysis.

Then the microbial features were preprocessed under three methods:

i. *original profile* with no further processing.
ii. *log-transformation:* all relative abundance values were transformed into logarithmic values.
iii. *batch-correction*: all relative abundance values were corrected by MMUPHin ^52^ R package (v.1.8.2) via the “adjust_batch” function to eliminate the batch effects, in which source study ID was set as the controlling factor.

## Meta-analysis

The beta diversity was measured by Permutational multivariate variance analysis (PERMANOVA; also referred to as Adnois test) that was implemented by package vegan ^58^ with 999 times permutation. *P*-value less than 0.01 denoted statistical significance. Principal coordinates analysis (PCoA) was performed by the labdsv package ^59^ based on the Bray-Curtis distance matrix. All meta-analysis was conducted using R software (version 4.1.3) ^60^.

We also used an ANOVA-like ^31^ analysis to quantify the effect of potential confounding factors and disease status on each microbial component. The total variance within the abundance of a given microbial feature was compared to the variance explained by disease status and the variance explained by confounders, based on an approach similar to linear models, including CRC status and various confounders (such as age, BMI, country, gender, and cohort origin) as explanatory variables. Variance calculations were performed on a rank basis to account for the non-Gaussian distribution of the microbiome abundance data ^31^. Continuous confounding variables were transformed into categorical variables either by quartile-based grouping or, in the case of BMI, by standard thresholds into normal weight (BMI ≤ 25), overweight (BMI 25–30), and obese (BMI > 30).

Plotting was performed using R (version 4.1.3), Python (version 3.9), CNSknowall (https://www.cnsknowall.com/), and Chiplot ^61^ (https://www.chiplot.online).

### Existing features selection tools

We compared seven feature selection tools using species-level microbiome profiles of the discovery dataset (**Table2**). Guided by the principle that methods yielding feature sets with greater overlap are likely to provide more reliable results ^62^, we first evaluated the consistency of CRC biomarkers selected by each tool. Among these, ANCOM-II was the most conservative, with 62% of its selected features overlapping with those identified by other methods. In contrast, RFECV detected the largest number of unique features, at 33%. Then based on the candidate biomarkers screened by each method, we employed XGBoost modeling with five-fold cross-validation to assess CRC detection performance on the discovery dataset. LEfSe achieved the highest AUROC, followed by RFECV and Wilcoxon test (**Fig. S6a**).

We also repeated the above test on each individual cohort of the ten discovery cohorts respectively. As shown in **Fig. S6b**, features selected by different tools varied widely in their distribution within single cohorts, measured by normalized proportion (NP). RFECV (mean NP: 38.36%) and LEfSe (mean NP: 11.44%) parsed out the highest rate, while ANCOM-II (mean NP: 2.4%), T-test (mean NP: 3.64%) and MaAsLin (mean NP: 4.05%) obtained the lowest NP (**Fig. S6b**). The XGBoost-based cross-validation (**Fig. S6c**) demonstrated that microbial markers identified by LEfSe significantly improved detection performance, exceeding the baseline accuracy of unfiltered profiles in almost all cohorts. Similarly, MaAsLin2 and Wilcoxon test also contributed largely to the accuracy of the XGBoost model. Although RFECV identified the largest number of candidate markers, its predictive performance was suboptimal likely due to the inclusion of false-positive features.

### Synergistic feature selection strategy

Considering the varying characteristics of the feature selection methods, here we proposed a synergistic strategy (**Fig. S6d**) to optimize biomarker screening while minimizing the impact of cohort-related variability. Specifically, we selected LEfSe, ANCOM-II, and MaAsLin2 for integration based on their complementary statistical foundations and demonstrated utility in microbiome studies. LEfSe combines non-parametric tests with linear discriminant analysis to detect biologically consistent features; ANCOM-II is tailored for zero-inflated and compositional data, reducing false positives in sparse microbiome matrices; and MaAsLin2 incorporates multivariable modeling to adjust for confounding factors such as age or BMI. These tools together offer a broad and balanced feature selection perspective across statistical paradigms. Firstly, LEfSe, ANCOME-II, and MaAsLin2 were applied to each single cohort and the all-merged data to identify a union set of candidate markers associated with CRC. Then this set was refined by the Max-Relevance and Min-Redundancy (mRMR) method ^67^ alongside an iterative feature elimination (IFE) ^48^ to finalize the optimal biomarkers for CRC.

### Machine learning models

*Model construction*. All machine learning-based (RF, XGBoost, MLP, KNN, and SVM) status detection was performed using the scikit-learn (version 1.3.2) package under Python 3.9. Among them, the neural network-based approach of MLP was implemented by PyTorch (version 1.11.0).

*Model evaluation*. Performance of model was evaluated using metrics including AUROC (area under the receiver operating characteristic curve), F1-score (*Eq. 1*), AUPRC (area under the precision-recall curve), and MCC (Matthews correlation coefficient; *Eq. 2*). Correlations among these metrics were measured by Pearson correlation coefficient.

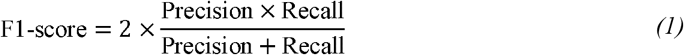

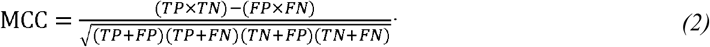

### Optimal workflow for CRC detection across multiple cohorts

To systematically evaluate the effects of preprocessing strategies, feature-selection methods, and classification algorithms on CRC prediction performance, we performed a comprehensive benchmarking analysis across ten discovery cohorts. This comprehensive analysis generated 1,080 testing combinations (**Fig. S7**). For each combination, the discovery dataset was randomly divided into training (70%) and test (30%) sets 100 times. Performance was assessed using AUROC, F1-score, AUPRC, and MCC, with AUROC selected as the primary metric due to its strong correlation with other measures (**Fig. S8**).

Based on the results of the benchmarked combinations (**Fig. S7**), no significant differences were observed in AUROC values among the three preprocessing strategies (two-sided Kruskal test, p-value = 0.4, **Fig. S9**). Considering logarithmic transformation reduced batch-related variability (**Fig. 2d**) with slightly better CRC detection effect than other pre-processed data (**Fig. S9**), subsequent analyses were conducted based on log-transformed profiles. In terms of annotation types, we observed that taxonomy profiles at the genus, species, and SGB levels outperformed other annotation types, yielding significantly higher AUROC values (two-sided Wilcoxon test, p-value < 0.01; **Fig. S7; Fig. S10a**), in agreement with confounder analysis results (**Fig. 2d**). For feature screening, biomarkers identified using the synergistic method reported slightly better AUROC values compared to individual tools (**Fig. S7** and **Fig. S10b**). Among machine learning models, XGBoost and RF significantly outperformed others in CRC prediction (two-sided Wilcoxon test, p-value < 0.01; **Fig. S7; Fig. S10c**), likely due to their capacity for handling high-dimensional, complex microbial data by non-linear modeling and noise resistance.

Building on these findings, we proposed an optimized bioinformatics workflow for microbiome-based CRC modeling (**Fig. 3a**), consisting of three key components:

i. *Preprocessing*. Annotate microbiome composition at the species level (with genus and SGB levels as optional alternatives) and apply log-transformation to relative abundance data to reduce skewness.
ii. *Feature selection*. Employ the synergistic feature approach to select disease-associated biomarkers.
iii. *Model construction*. Construct and adjust the model using RF (XGBoost as an alternative).

### Instance-based transfer learning for cross-cohort ADA detection

i. *Subset selection and matching by a mutual nearest neighbor set algorithm*: Randomly select a subset (e.g., 30% samples) *T’* from the target domain *T*. For each sample *t’*_*i*_ in *T’*, we search for its top *k* (e.g., k=10) nearest neighbors from source domain *S* by Euclidean distance, denoting as set *KNN*_*Euclidean*_ (*S, t’*_*i*_). Similarly, for each sample *s*_*i*_ in the source domain *S*, we also search its *KNN*_*Euclidean*_ (*T’, s*_*i*_) set from *T’*. Here we define a sample pair *s*_*i*_ ∈ S and *t’*_*j*_ ∈ *T’* as pre-matched neighbor samples *N*_*Euclidean*_ (*s*_*i*_, *t’*_*j*_) if and only if under *Eq. 3*.

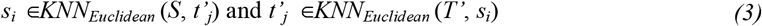

Then such pairing relation was re-confirmed by Cosine distance to determine the final matches *N*_*Cosine*_ (*s*_*i*_, *t’*_*j*_). Finally, according to *Eq. 4*, we take the intersection of *N*_*Euclidean*_ (*si, t’j*) and *N*_*Cosine*_ (*s*_*i*_, *t’*_*j*_), thereby forming the mutual nearest neighbor set (MNN).

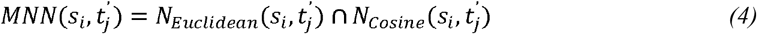
ii. *Feature extraction and initial model*: Utilize matched sample pairs *MNN* (*s*_*i*_, *t’*_*j*_) from the source and target domains to identify ADA-relevant features through synergistic feature selection. Subsequently, construct an initial Random Forest (RF) model using *MNN* (*s*_*i*_) from the source domain.
iii. *Pseudo labeling*: Apply the initial model to parse out the detected status (*i*.*e*., assign pseudo-labels) of each sample in target subset *T’*, forming a new set as *T’’*.
iv. *Dataset fusion*: Combining *T’’* with pseudo labels with the target domain *T’* to create an enhanced dataset *E (i*.*e*., *E* = *T’’* + *T’*).
v. *Final model training*: Train the final TL model on the enhanced dataset *E* using the optimal bioinformatics workflow (**Fig. S10**).

### Jensen-Shannon divergence analysis

To evaluate the distributional similarity between the source and target domains before and after transfer learning, we first normalize the feature values of each sample (*Eq. 5*).

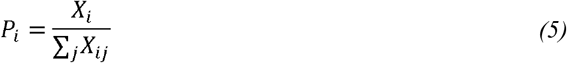

Here, *X*_*i*_ represents the feature values of the *i*-th sample, and *X*_*ij*_ represents the value of the *j*-th feature for the *i*-th sample. Subsequently, we employ the Jensen-Shannon (JS) divergence (*Eq. 6*) to assess the similarity between the distributions of the two datasets. Each sample in the datasets was represented as a probability distribution over the feature space.

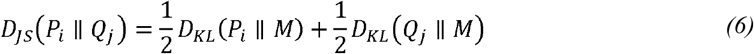

Here, *P*_*i*_ and *Q*_*j*_ represent the probability distributions of the *i*-th sample from the target domain and the *j*-th sample from the source domain, respectively. where *M* = (*P*_*i*_ + *Q*_*j*_)/2 is the mean distribution and *D*_*KL*_ (*P* || *M*) is the Kullback-Leibler (KL) divergence (*Eq. 7)*, which forms the basis of the JS divergence, quantifies the information loss when approximating one distribution *P* with another *Q*.

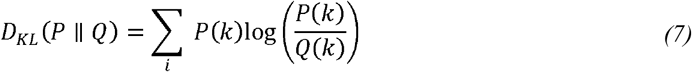

Here, represent the features in the probability distribution of a single sample, such that *P(k)* and *Q*(*k*) denote the probabilities at the *k*-th feature for the respective samples.

The JS divergence is symmetric, satisfying *D*_*JS*_ (*P*_*i*_ || *Q*_*j*_) = *D*_*JS*_ (*Q*_*j*_ || *P*_*i*_) and ranges between 0 and 1, where smaller values indicate greater similarity between the distributions. Thus, smaller JS divergence values suggest higher similarity between the sample distributions.

## Data availability

The in-house produced (CHN_WF cohort) dataset has been deposited at the Sequence Read Archive (SRA) with project ID PRJNA1198725. The public datasets are available at SRA and ENA via identifiers listed in **Table 1**.

The source datasets are available at the GitHub repository including:

## Supplementary tables

https://github.com/qdu-bioinfo/CRC_project/blob/main/Supplementary/CRC-Supplementary_tables.xlsx

**Dataset S1:**

https://github.com/qdu-bioinfo/CRC_project/blob/main/Supplementary/CRC-Dataset_S1.xlsx

**Dataset S2:**

https://github.com/qdu-bioinfo/CRC_project/blob/main/Supplementary/CRC-Dataset_S2.xlsx

## Code availability

Source codes and analysis scripts are available at the GitHub repository at https://github.com/qdu-bioinfo/CRC_project.

## Authors’ contribution

X.S. and X.F. conceived and designed the project. Y.S. and J.Z. designed the algorithms and workflow. X.F. collected the gut microbiome samples. Y.S., Z.W., J.X. and S.W. collected the public datasets. Y.S., W.Z. and H.G. performed the data analysis. X.S. and Y.S. drafted the manuscript. All authors read and approved the final manuscript.

## Acknowledgements

X.S. acknowledges support of grant 2021YFF0704500 from National Key Research and Development Program of China, grant 32070086 from National Natural Science Foundation of China, Shandong Province Youth Entrepreneurial Talent Introduction and Training Program, and Shandong Province Taishan Scholars Youth Experts Program. All authors thank Mr. Yi Zhao from Qingdao University for the support of computing resources.

