## Supplementary figures for "Instance-based Transfer Learning Enables Cross-Cohort Early Detection of Colorectal Cancer"

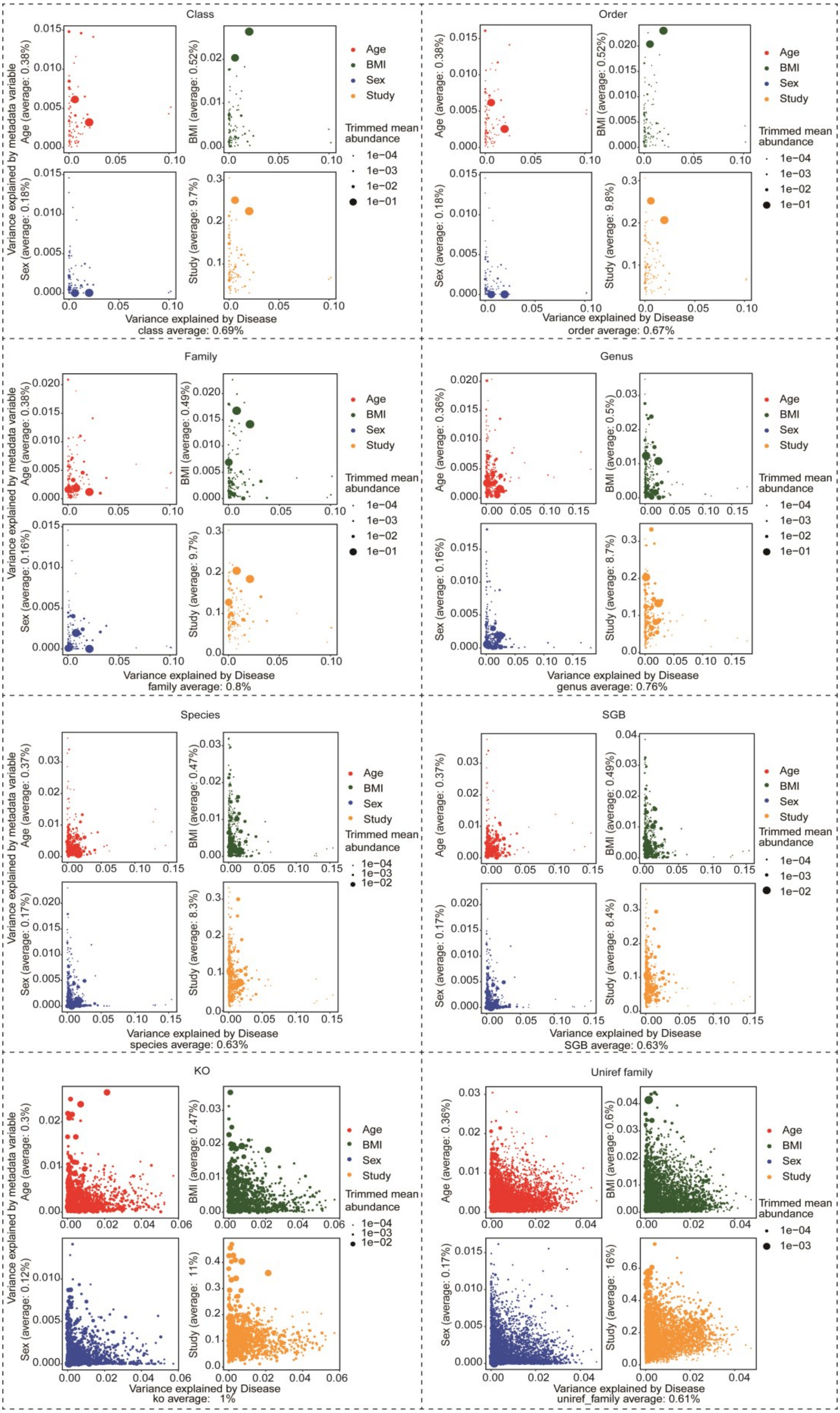

**Fig. S1. Variance in microbial composition attributed to health status and other confounding factors.** The variance explained by disease and the variance explained by other factors (age, BMI, disease status, gender, and cohort) are plotted in the same coordinate system. The horizontal axis represents the amount of variance explained by disease, and the vertical axis represents the amount of variance explained by confounders for microbial composition. The abundance of each taxon is represented by the size of the point. Source data is available in Dataset S2.

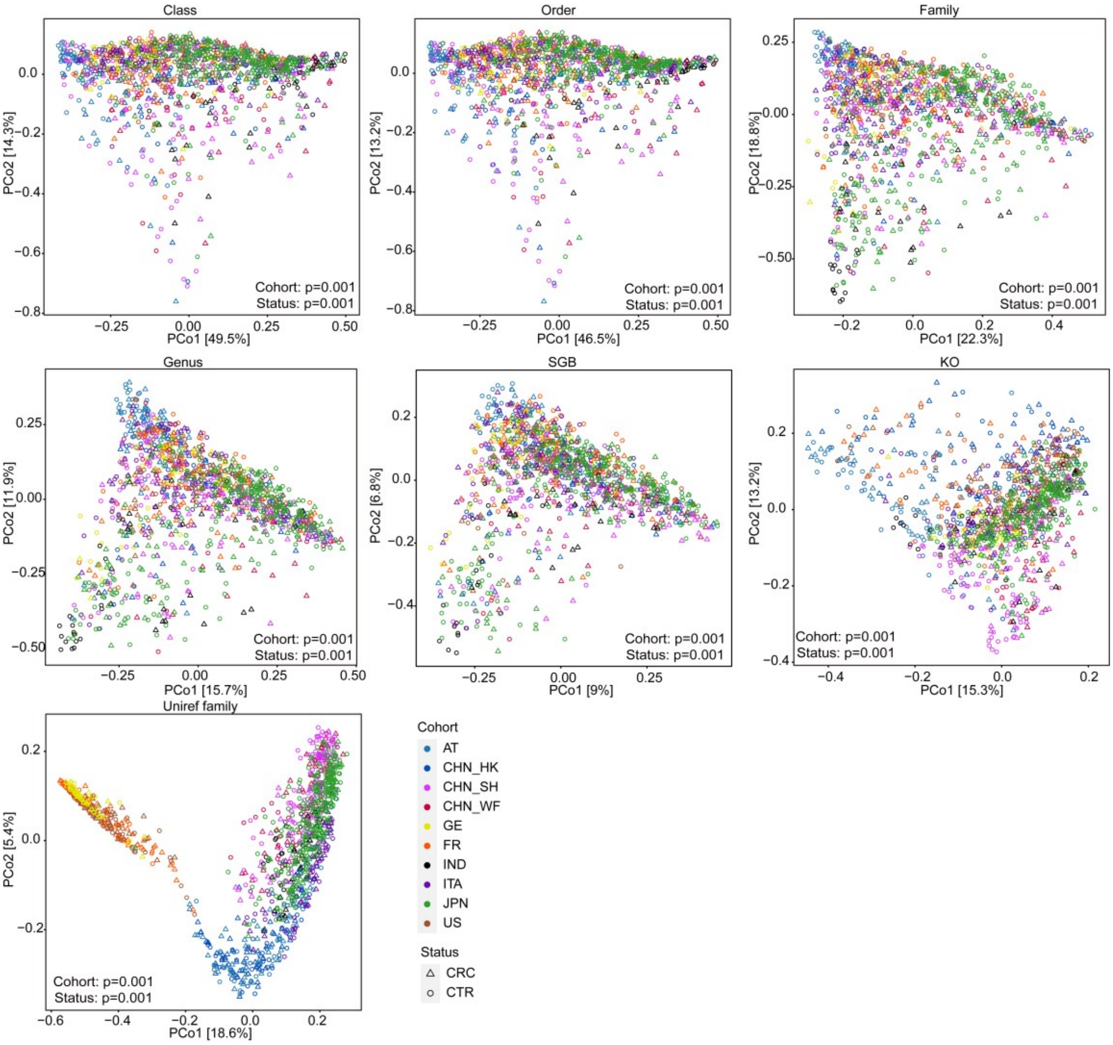


**Fig. S2. Beta diversity patterns of CRC and CTR samples in the discovery dataset, visualized using Bray-Curtis distances at seven annotation levels.** Statistical significance was assessed by the PERMANOVA test for beta diversity, with corresponding *p*-values reported.

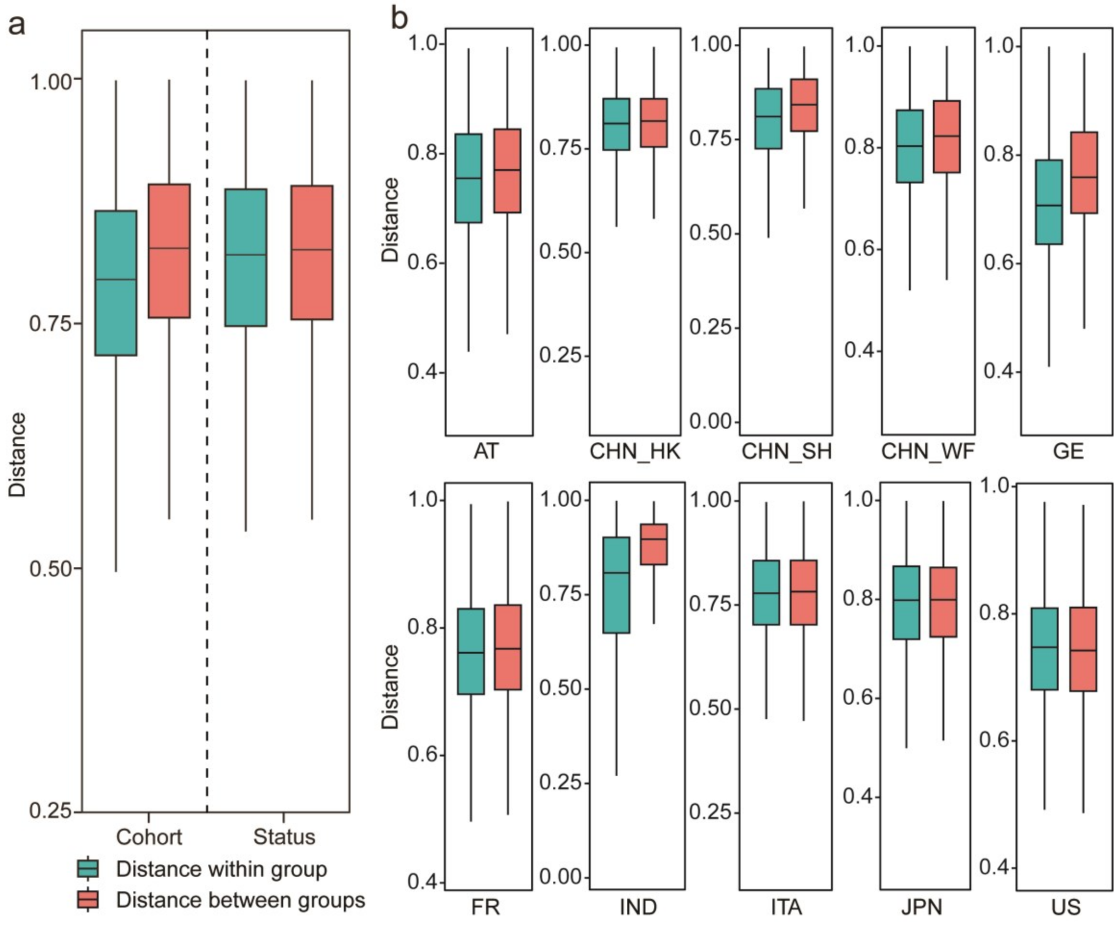


**Fig. S3. Bray-Curtis distances across disease states and cohorts on species level.** (a) Distances of samples in different health states and cohorts. (b) Distances of samples in different health states in each single cohort.

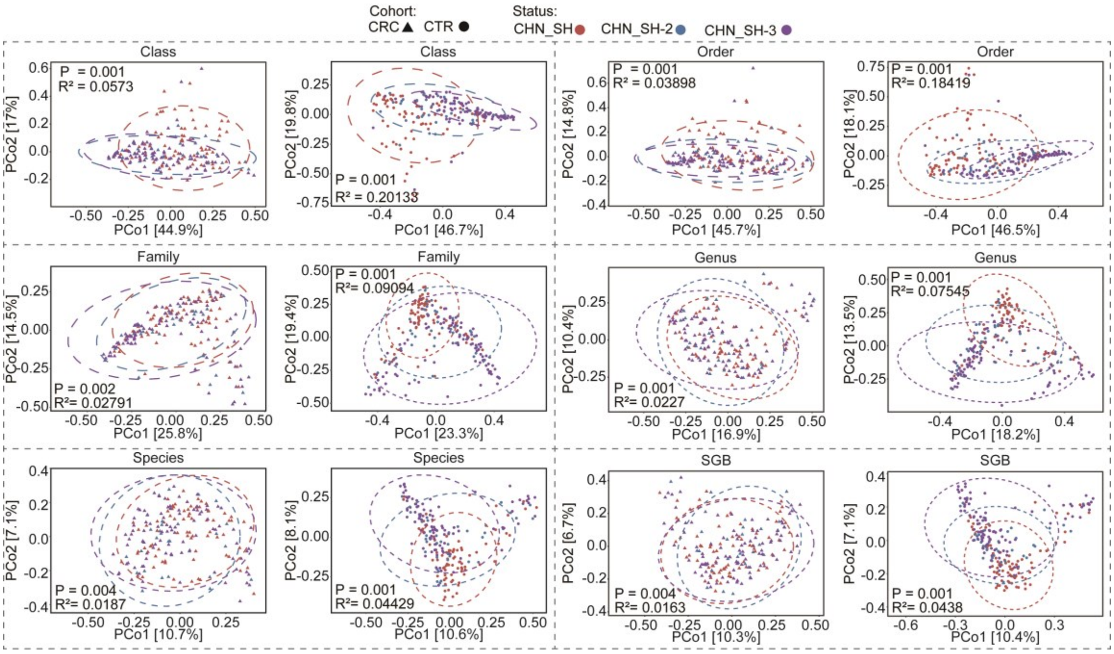


**Fig. S4. Technical batch effect on gut microbial composition**. Principal coordinate analysis (PCoA) was performed on three cohorts (CHN_SH, CHN_SH-2, CHN_SH-3) collected from the same region by six types of microbial features. Triangles represent colorectal cancer (CRC) samples, circles represent healthy (CTR) samples, and different colors represent samples from different cohorts. Statistical significance was assessed by the PERMANOVA test for beta diversity, with corresponding *p*-values reported.

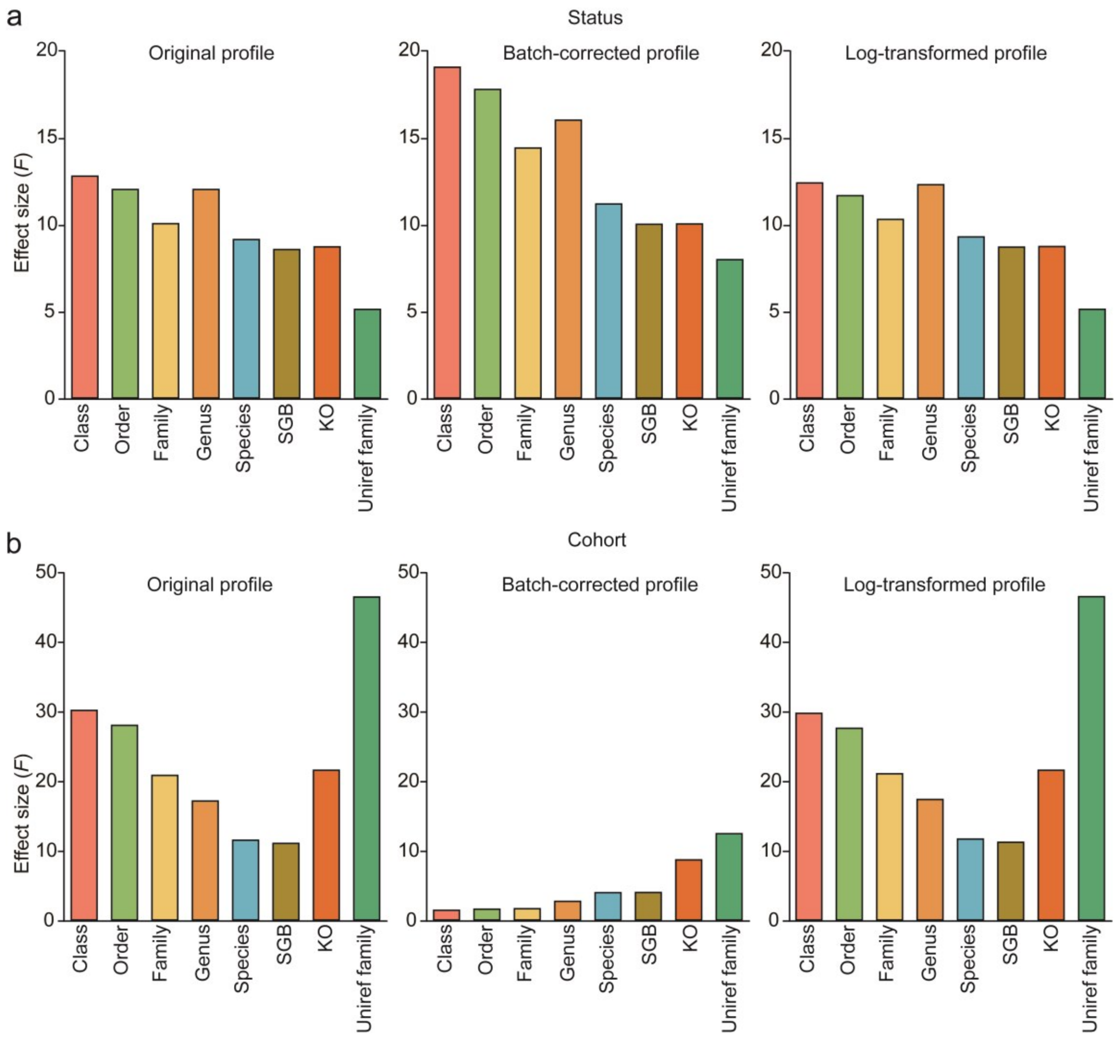


**Fig. S5. Effect sizes of health status (a) and cohort (b) across preprocessing strategies and microbial feature types.** *F*-values were derived from the PERMANOVA test. Source data is available in Dataset S2.

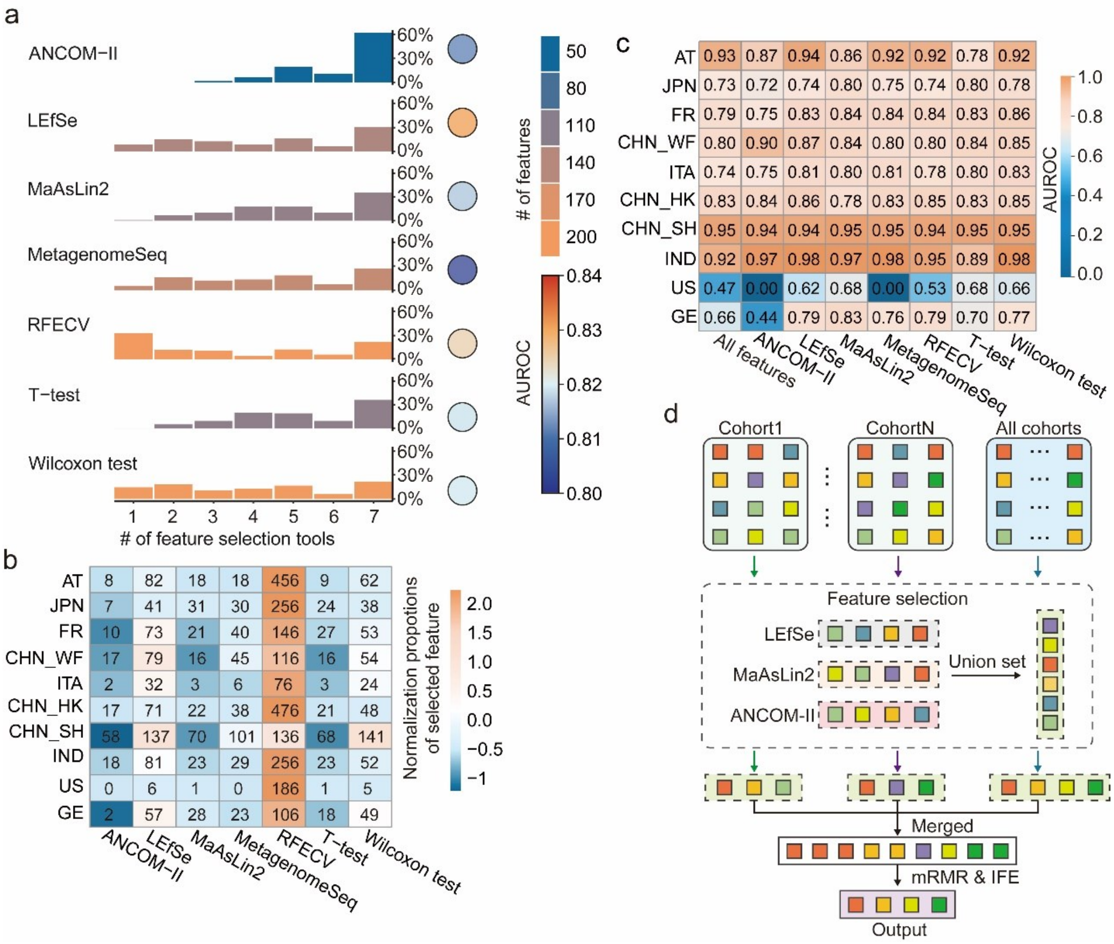


**Fig. S6. Comparison of different feature selection tools.** (a) Performance of different tools on all cohorts of the discovery dataset. For each method, the color of the bars indicates the number of identified features shared with others, and the height represents their proportion within a method. The colors of the dot chart illustrated the AUROC of the XGBoost-based cross-validation using features selected by each method. (b) Feature selection of different methods on each single cohort of the discovery dataset. Values are the number of identified microbial markers, and colors are their normalized proportion (NP) by scaled mean-centralization in each single cohort (each row). (c) Performance of XGBoost-based cross-validation in each single cohort. (d) Diagram of the synergistic feature selection strategy with mRMR and IFE.

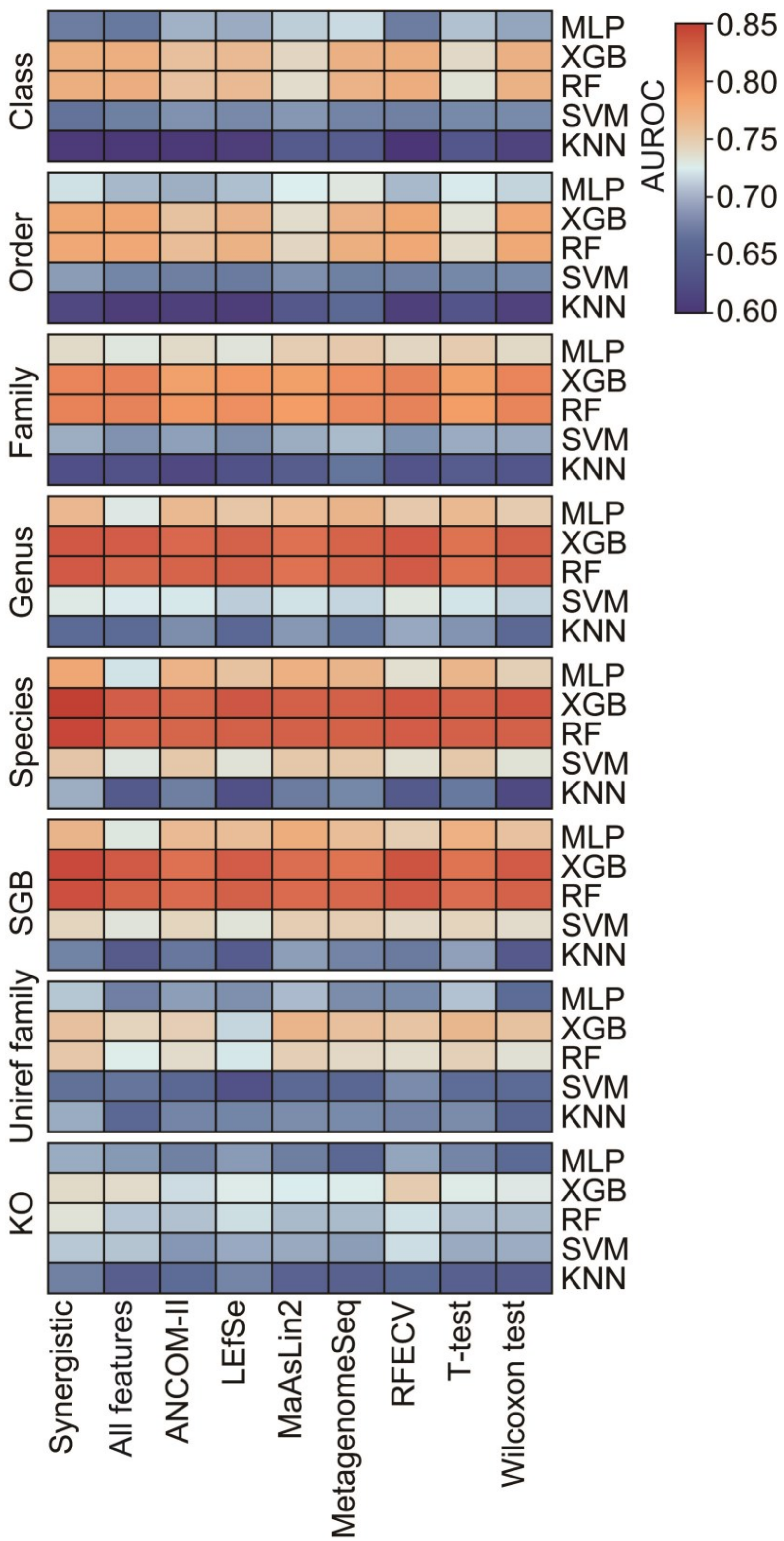


**Fig. S7. Comprehensive AUROC analysis for CRC detection.** The heatmap displays 360 AUROC results by five machine learning, nine feature selection methods and eight feature types using logarithmically transformed profiles. Results of all 1,080 test combinations are available in Dataset S2.

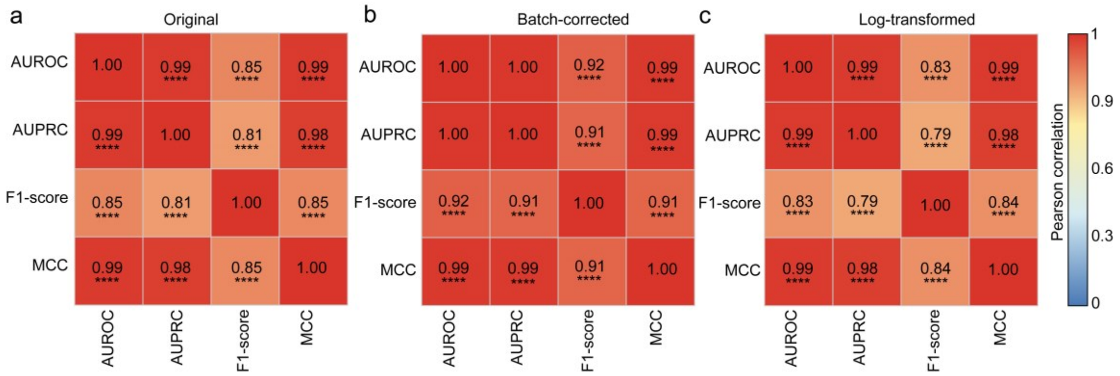


**Fig. S8. Correlation analysis of evaluation metrics.** Heatmaps illustrate the pairwise correlation analysis results among four evaluation metrics: AUROC, AUPRC, F1-score, and MCC. This analysis was performed using Pearson correlation on microbial profiles with three preprocessing strategies: (a) original data, (b) batch-corrected data, and (c) logarithm-transformed data. Significance is indicated as: * for $p$-value < 0.05, ** for $p$-value < 0.01, *** for $p$-value < 0.005, and **** for $p$-value < 0.001. Source data is available in Dataset S2.

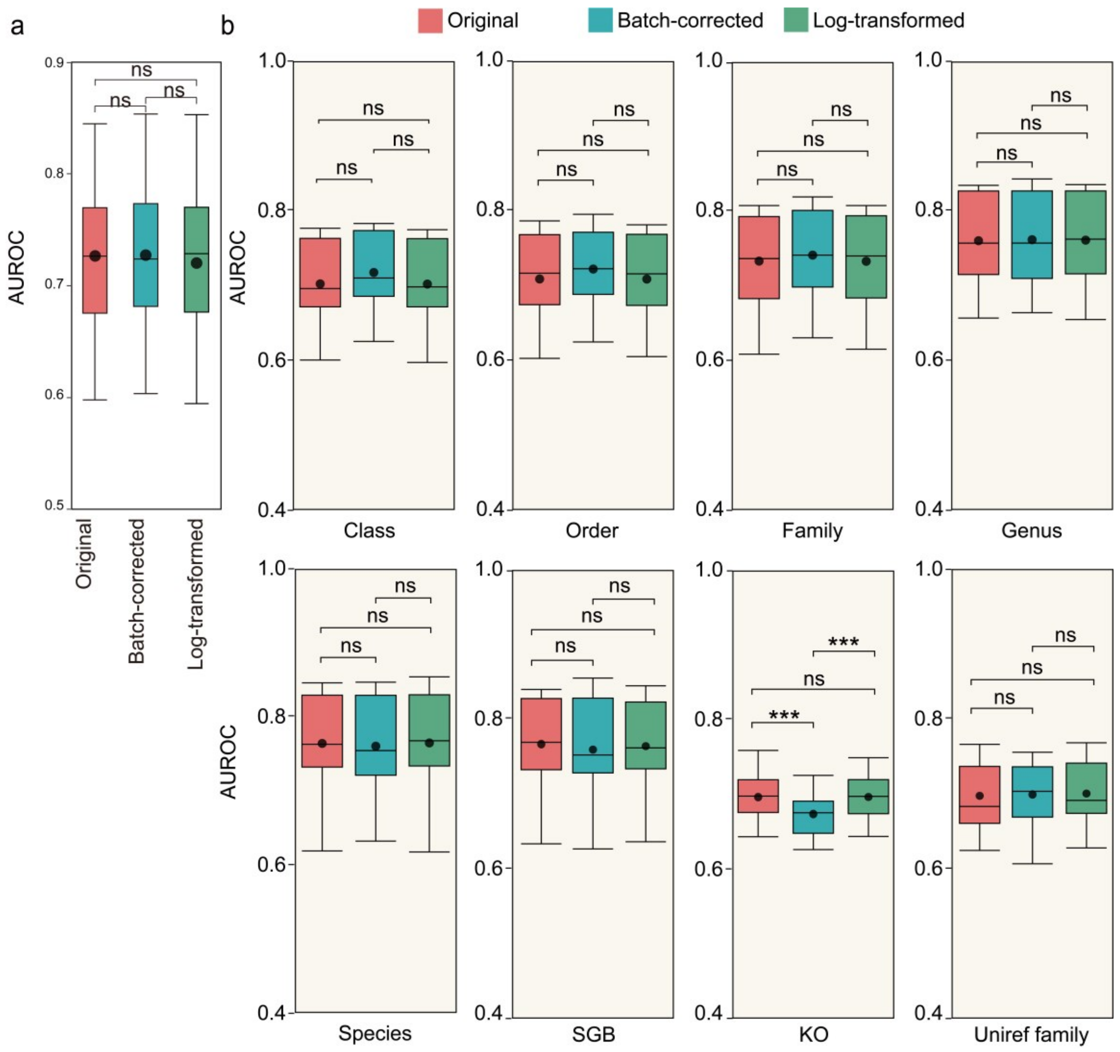


**Fig. S9. Comparison of AUROC across preprocessing strategies.** (a) Difference of AUROC among preprocessing strategies measured by two-sided Wilcoxon test. (b) Statistical analysis was performed using the two-sided Wilcoxon test. Significance is indicated as: * for *p*-value < 0.05, ** for *p*-value<0.01, *** for *p*-value<0.001, and ns for *p*-value > 0.05. Source data is available in Dataset S2.

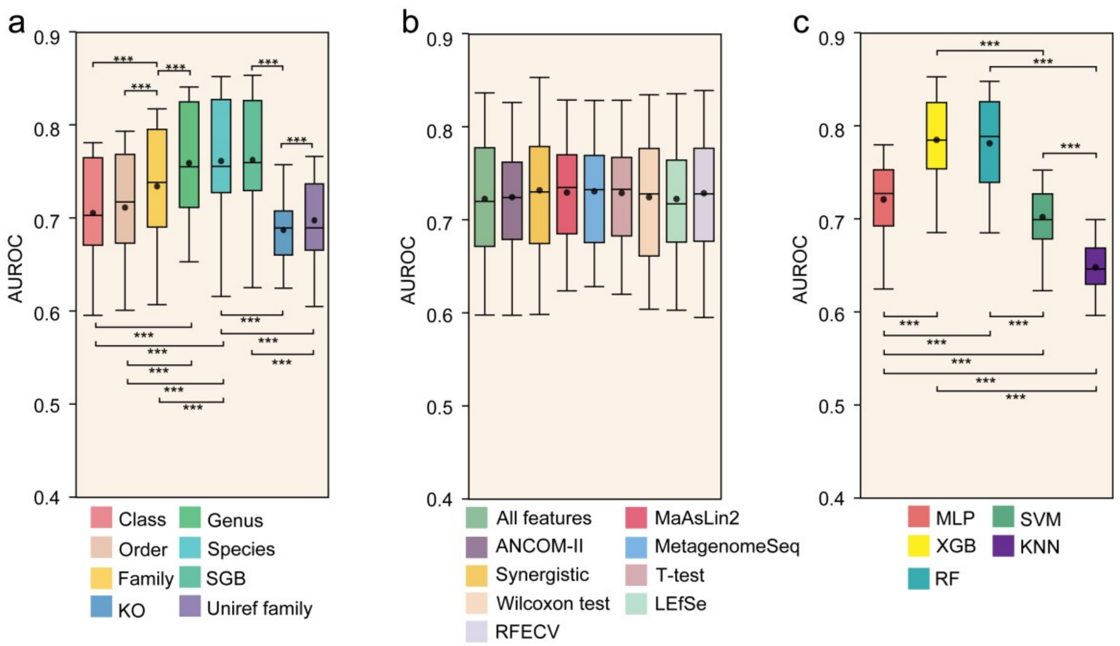


**Fig. S10. Performance variation of each single step for CRC detection.** (a) AUROC results of detecting CRC by different feature types. (b) AUROC results of CRC detection using various feature selection tools. (c) AUROC results for CRC detection using different machine learning models. Statistical analysis was performed using the two-sided Wilcoxon test. Significance is indicated as: * for p-value < 0.05, ** for p-value<0.01, and *** for p-value<0.001. Source data is available in Dataset S2.

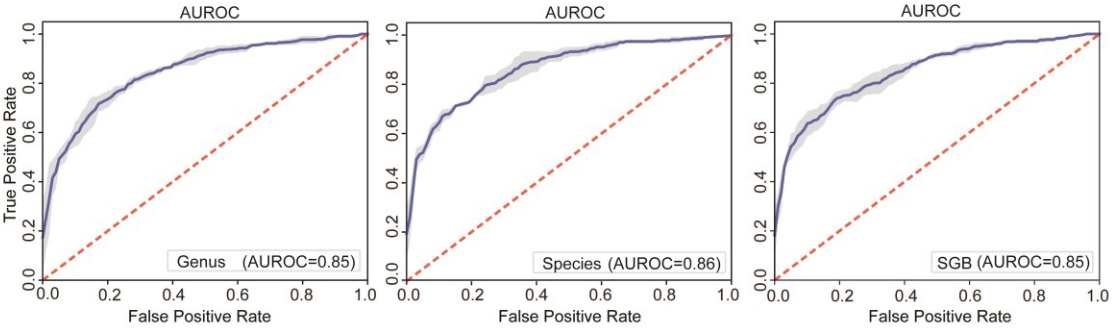


**Fig. S11**. **AUROC of the optimal workflow using Random Forest model on three suggested types of microbial features.** Source data is available in Dataset S2.

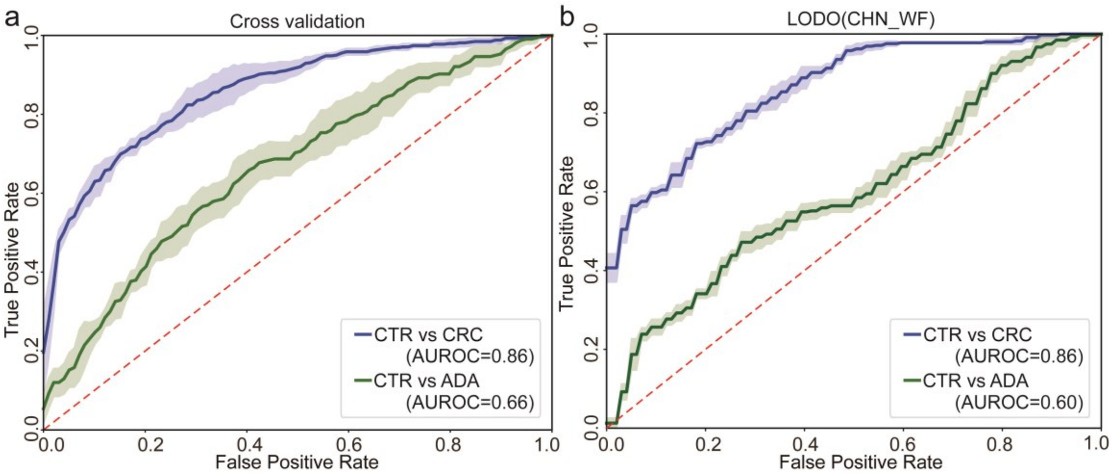


**Fig. S12. Detecting ADA across cohorts.** (a) ROC curves of CRC and ADA detection on multiple cohorts by five-fold cross validation. (b) ROC curves of CRC and ADA detection on CHN_WF cohort by other cohorts. Source data is available in Dataset S2.

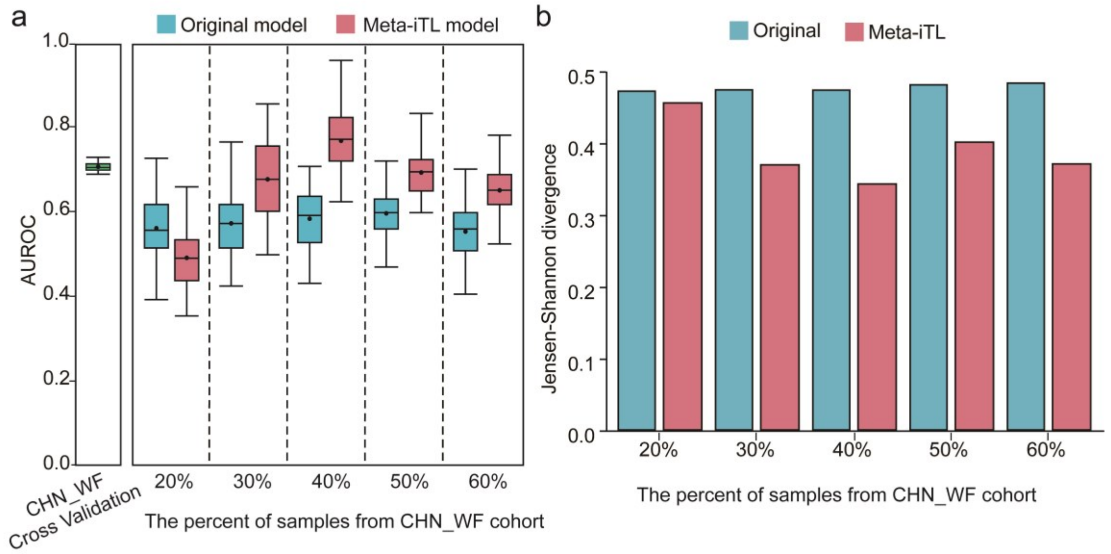


**Fig. S13. Meta-iTL significantly improves the performance for ADA detection across cohorts.** (a) Comparison of performance of ADA detection for target domain (CHN_WF cohort) using model trained by source domain (all other cohorts) before and after TL. The baseline is a five-fold cross-validation within the target domain. (b) Jensen-Shannon (JS) values between the data before and after TL and the target domain. Source data is available in Dataset S2.

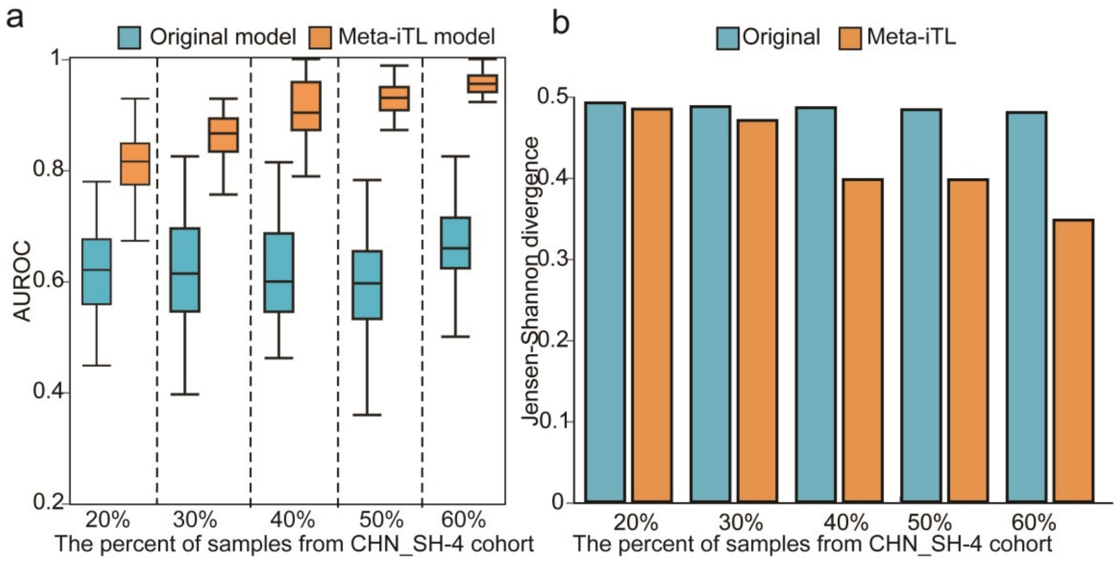


**Fig. S14. Meta-iTL significantly improves the performance of ADA detection across cohorts**. (a) Comparison of performance of ADA detection for target domain (CHN_SH-4 cohort) using model trained by source domain (all other cohorts) before and after TL. The baseline is a five-fold cross-validation within the target domain. (b) Jensen-Shannon (JS) values between the data before and after TL and the target domain. Source data is available in Dataset S2.
